# The immunodominant and neutralization linear epitopes for SARS-CoV-2

**DOI:** 10.1101/2020.08.27.267716

**Authors:** Shuai Lu, Xi-xiu Xie, Lei Zhao, Bin Wang, Jie Zhu, Ting-rui Yang, Guang-wen Yang, Mei Ji, Cui-ping Lv, Jian Xue, Er-hei Dai, Xi-ming Fu, Dong-qun Liu, Lun zhang, Sheng-jie Hou, Xiao-lin Yu, Yu-ling Wang, Hui-xia Gao, Xue-han Shi, Chang-wen Ke, Bi-xia Ke, Chun-guo Jiang, Rui-tian Liu

## Abstract

The coronavirus disease 2019 (COVID-19) pandemic caused by severe acute respiratory syndrome coronavirus 2 (SARS-CoV-2) becomes a tremendous threat to global health. Although vaccines against the virus are under development, the antigen epitopes on the virus and their immunogenicity are poorly understood. Here, we simulated the three-dimensional structures of SARS-CoV-2 proteins with high performance computer, predicted the B cell epitopes on spike (S), envelope (E), membrane (M), and nucleocapsid (N) proteins of SARS-CoV-2 using structure-based approaches, and then validated the epitope immunogenicity by immunizing mice. Almost all 33 predicted epitopes effectively induced antibody production, six of which were immunodominant epitopes in patients identified via the binding of epitopes with the sera from domestic and imported COVID-19 patients, and 23 were conserved within SARS-CoV-2, SARS-CoV and bat coronavirus RaTG13. We also found that the immunodominant epitopes of domestic SARS-CoV-2 were different from that of the imported, which may be caused by the mutations on S (G614D) and N proteins. Importantly, we validated that eight epitopes on S protein elicited neutralizing antibodies that blocked the cell entry of both D614 and G614 pseudo-virus of SARS-CoV-2, three and nine epitopes induced D614 or G614 neutralizing antibodies, respectively. Our present study shed light on the immunodominance, neutralization, and conserved epitopes on SARS-CoV-2 which are potently used for the diagnosis, virus classification and the vaccine design tackling inefficiency, virus mutation and different species of coronaviruses.

## INTRODUCTION

The coronavirus disease 2019 (COVID-19) pandemic caused by the novel severe acute respiratory syndrome coronavirus 2 (SARS-CoV-2) has caused unprecedented impact on global health. More than 6 million cases were reported by WHO on June 1, 2020 (https://covid19.who.int/) (Guan et al., 2020; Zhu et al., 2020). SARS-CoV-2 shares 96.2% and 79.5% genomic sequence identity with bat coronavirus and SARS-CoV, respectively, but it is more contagious than SARS-CoV (Lu et al., 2020; Zhou et al., 2020). Prophylactic vaccines are an important means to curb the pandemic of infectious diseases. Accordingly, effective and safe SARS-CoV-2 vaccine is urgently needed. Eight SARS-CoV-2 vaccine candidates based on a variety of technologies are being tested in clinical trials (Chen et al., 2020a; Thanh Le et al., 2020). However, the epitopes on these vaccines and SARS-CoV-2 are not well-studied, and it is still urgent to identify epitopes that can elicit neutralizing antibodies and determine the immunodominant epitopes in humans for the improvement and design of novel vaccines.

Four major structural proteins, spike (S), envelope (E), membrane (M), and nucleocapsid (N) proteins play vital roles in entry and replication of the virus (Chen et al., 2020b). Several epitopes on S protein have been reported although with little information of the immunogenicity and the neutralization, but the most epitopes on M, E, and N proteins still remain unknown (Baruah and Bose, 2020; Bhattacharya et al., 2020; Yuan et al., 2020a, b). The accuracy of the predicted epitopes using *in silico* methods is unclear and the immunogenicity of the obtained epitopes needs further experimental verification (Ahmed et al., 2020; Grifoni et al., 2020; Kiyotani et al., 2020). Epitope prediction methods based on the three-dimensional structure of protein can greatly improve the precision of antigen epitopes (Jespersen et al., 2017). Therefore, the reported 3D structures of S and M proteins are conductive to epitope prediction (Jin et al., 2020; Lan et al., 2020; Walls et al., 2020; Wrapp et al., 2020). Although the structures of E and N proteins are still unsolved, it is possible to model these protein structures based on their reported gene sequence using molecular simulation and then predict their epitopes (Lu et al., 2020). Increasing evidences showed that some linear epitopes, as the sites of virus vulnerability, conserved regions or the key components of conformational epitopes, play important roles in the induction of virus neutralization (Alphs et al., 2008; Sok and Burton, 2018; Xu et al., 2018). For example, a linear epitope of HIV induced broad-spectrum protection effect, and could be used to develop universal vaccines (Kong et al., 2019). By identifying the conformational B cell epitopes with higher degree of continuity and the appropriate linear window with key functional residues of discontinuous B cell epitopes centralized and randomized, we may find the key components of conformational epitopes.

The immunogenicity, immunodominance, especially neutralization of the epitopes is crucial for the development of effective SARS-CoV-2 vaccines. Although the epitope immunodominance landscape of S protein was mapped (Zhang et al., 2020), mutation on virus proteins might alter the antigenicity of the virus and possibly affected human immune responses to the epitopes, making it the central challenge for the vaccine development. Phylogenetic analysis showed that SARS-CoV-2 mutated with a mutation rate around 1.8 × 10^−3^ substitutions per site per year (Li et al., 2020). Within all the identified mutations of S protein, further investigation is needed on the 614th amino acid. G614 in S1 protein of SARS-CoV-2, found in almost all the COVID-19 patients outside China, exhibited higher case fatality rate and might be more easily spread than D614 which mainly found in China (Becerra-Flores and Cardozo, 2020). The 614th amino acid is located on the surface of S protein protomer and the G614 destabilized the conformation of viral spike and facilitated the binding of S protein to ACE2 on human host cells (Becerra-Flores and Cardozo, 2020). However, little is known about how G614 influences human immune responses to SARS-CoV-2. In fact, the mutations not only on S protein, but also on E, M, N proteins might affect human immune responses to the virus. The limited neutralizing effect by the vaccine using S protein as the only antigen suggested that epitopes on E, M, and N proteins might be important for SARS-CoV-2 vaccine design as well, and understanding how mutations affect human immune responses to the virus is necessary (van Doremalen et al., 2020).

In this study, we predicted and synthesized the B cell epitopes on the surface of S, M, E, N proteins of SARS-CoV-2, prepared 37 vaccines based on HBc virus-like particles (VLP) using SpyCatcher/SpyTag system, validated the immunogenicity of the epitopes by immunizing mice, and identified epitopes that could elicit neutralizing antibodies. We also determined immunodominant epitopes on SARS-CoV-2 by mapping the epitopes with the sera from COVID-19 convalescent patients and analyzed the relevance between epitope immunodominance and the mutations on SARS-CoV-2 proteins.

## RESULTS

### Prediction of SARS-CoV-2 B cell epitopes and preparation of HBc-S VLPs displayed with the epitopes

In order to predict antigen epitopes on SARS-CoV-2, we used high performance computer to simulate the three-dimensional structures of S, M, E, N proteins, and then used computational simulation calculations to obtain preliminary antigen epitope information based on epitope surface accessibility. The spatial structure information of S, M, E, and N protein structure models was obtained (Fig. 1A-D), in which structures of S and M were consistent with the reported structures and the structures of E and N were obtained for the first time (Wrapp et al., 2020; Yan et al., 2020). A total of 33 B-cell epitopes were predicted on the basis of protein structures and the priority was given during the selection process to select sequences with high homology within SARS-CoV and RaTG13 coronavirus strains (Table S1, Fig. S1). Within these epitopes, four of them contained glycosylation sites (Watanabe et al., 2020; Wrapp et al., 2020) and the according GlcNAc glycosylated epitopes were synthesized (Table S1), and 13 from S protein, 2 from E protein, 3 from M protein, and 5 from N protein, respectively, are conserved with >80% homology among SARS-CoV-2, SARS-CoV and bat coronavirus RaTG13 (Table S1). All the epitopes were exposed on the surface of the virus (Fig. 1A-D) and had a high antigenicity score, indicating their potentials in initiating immune responses. Therefore, they were considered to be promising epitope candidates against B-cells for vaccine preparation.

**Figure 1.**
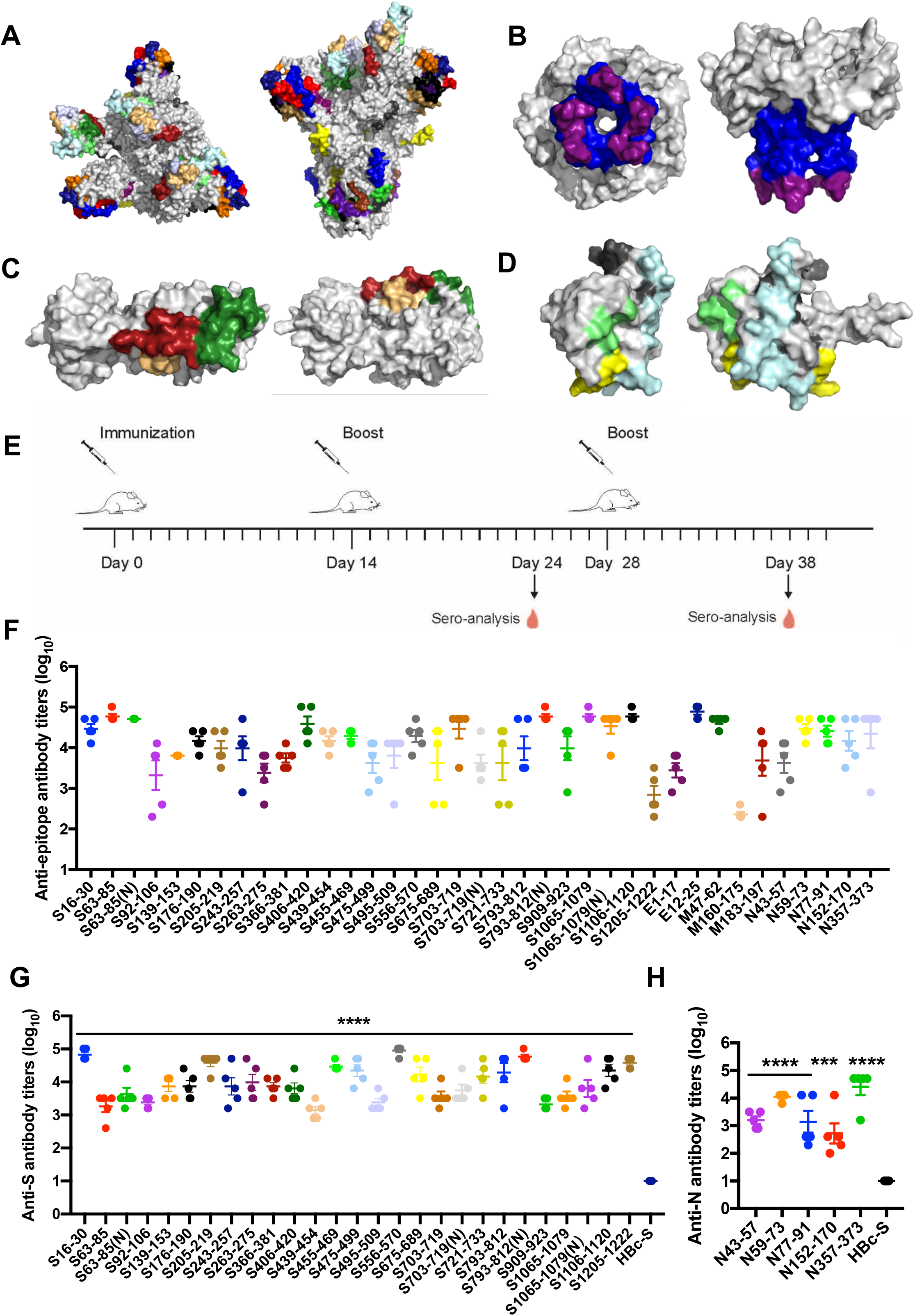
Predication and validation of epitopes on SARS-CoV-2. (A-D) Molecular simulated structures and predicted epitopes of major proteins of SARS-CoV-2. Top and side views of three-dimensional structures (grey) and the predicted epitopes (colored) of spike protein (S) (A), envelope protein (E) (B), membrane protein (M) (C), and nucleocapsid protein (N) (D). (E-G) Epitope conjugated HBc-S VLPs induce high antibody titers against epitope peptides and SARS-CoV-2 proteins. (E) Immunization schematic design. BALB/c mice (female, 6-8 weeks, n=5) were immunized with HBc-S-P VLPs for 3 times, respectively. (F-H) 96-well plates were coated with peptides (F), S (G) or N (H) proteins, respectively. The sera from mice immunized by HBc-S decorated with epitopes from S, M, E, N proteins were serially diluted from 1:100 to 1:102400 in two-fold dilution steps and added to the plates. Results are shown as mean ± SEM (Compared with HBc-S control; ***p < 0.001; ****p < 0.0001; one-way ANOVA followed by Dunnett’s test; Compared with non-glycosylated epitope; #p < 0.05; Student t-test).

The predicted epitope peptides of S, M, E, and N proteins were synthesized and conjugated onto the surface of HBc-S VLPs via SpyCatcher/SpyTag isopeptide formation, respectively, forming epitope peptide displaying HBc-S VLPs, termed as HBc-S-P VLPs. SDS-PAGE confirmed that the epitope peptides were successfully conjugated onto HBc-S VLPs shown by HBc monomers with higher molecular weight (Fig. S2A). Our TEM results showed that all the HBc-S-P self-assembled into morphologically correct VLPs (Fig. S2B). We further assessed the hydrodynamic diameter of HBc-S-P VLPs by DLS, and the results showed that all the HBc-S-P VLPs were relatively uniform (PDI < 0.25) with a diameter around 40 nm (Fig. S2C, Table S2), which was consistent with previous reports (Ji et al., 2018).

### The epitopes elicit highly specific antibody responses

To assess the immunogenicity, the HBc-S-P VLPs were subcutaneously immunized to BALB/c mice for three times and the serum antibody titers were assayed by ELISA (Fig. 1E). Epitopes S455-469, S556-570, E12-25, M47-62, N59-73, and N357-373 elicited robust antibody responses against peptides and/or S, N proteins in mice at 10 days after the second injection (≥1000, Fig. S3). All the predicted epitopes boosted antibodies response against the corresponding epitope peptides and the antibody titer reached at least 1000 after the third immunization, except for M160-170 with antibody titer being only 230 (Fig. 1F). Accordingly, the epitopes S63-85, S205-219, S455-469, S475-499, S556-570, S721-733, S793-812, S1106-1120, S1205-1222, N59-73, N353-373 on S and N proteins also induced robust antibodies with titers greater than 10000 against S and N proteins, respectively (Fig. 1G and 1H). These results demonstrated that almost all the predicted epitopes on S, M, E, and N proteins elicited immune responses with high levels of antibodies, suggesting these epitopes have good immunogenicity. The GlcNAc glycosylated epitopes also elicited sufficient amount of antibodies towards the corresponding epitope peptides and S protein, respectively, and only S793-812(N) induced higher antibodies than that of the non-glycosylated epitope (Fig. 1F and 1G).

### Imported and domestic COVID-19 cases have different immunodominant epitopes

To investigate the spectrum of antibodies in COVID-19 patients, we detected the binding of the early convalescent sera of 8 imported (Europe) cases which infected SARS-CoV-2 in early April, 2020 and 12 domestic (China) cases in early February, 2020 to various epitopes (Table 1). The mean value plus three times of the standard deviation in healthy volunteers was used as the cut-off value to define positive reactions and the epitope showing the average positive rate ≥50% among patients was considered as an immunodominant epitope (Fig. 2A and S4). Our results showed that S556-570, M183-197 and N357-373 were immunodominant in domestic COVID-19 patients (Fig. 2B-D), whereas S675-689 and S721-733 were immunodominant in imported COVID-19 groups (Fig. 2E-F). Only N152-170 was immunodominant in both groups (Fig. 2G). Notably, S556-570, N152-170 and S721-733 reacted with the sera of almost all the patients (≥80%) in domestic or imported cases, respectively (Fig. 2B, 2F and 2G). These results indicated that imported and domestic COVID-19 cases had different immunodominant epitopes.

**Figure 2.**
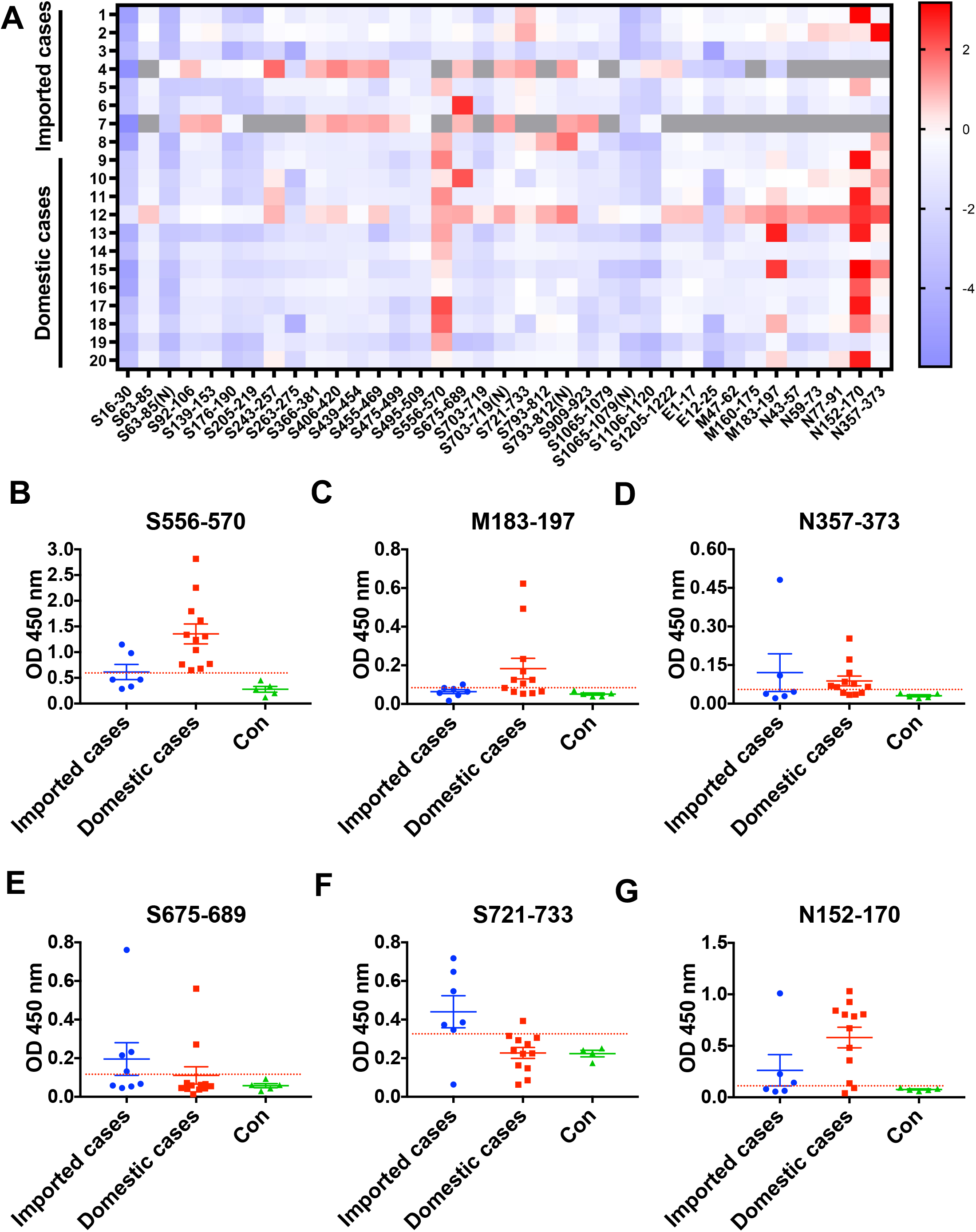
Imported and domestic COVID-19 cases have different immunodominant epitopes. (A) The landscape of adjusted epitope-specific antibody levels in early convalescent sera of imported and domestic COVID-19 patients. The ELISA results of absorbance at 450 nm were normalized to the aforementioned cut-off values. Gray indicated not tested. (B-G) Immunodominant epitopes binding with the antibodies in early convalescent sera from imported and domestic COVID-19 patients. 96-well plates were coated with 0.5 μg peptides and sera were diluted at 1:50. The cut-off lines were based on the mean value plus 3SD in 4-5 healthy persons.

To elucidate the possible cause of the difference in immunodominant epitopes, we sequenced the S, M, N, and E genes of SARS-CoV-2 from imported and domestic COVID-19 patients. The results showed that the sequences of E and M proteins were identical in both imported and domestic COVID-19 patients. However, the gene sequences of imported and domestic strains contained G614 or D614 in S protein, and K203R204/G189R203G204/R203G204/R203G204S344 in N protein, respectively (Table 1), resulting in different immunodominant epitopes of different virus sub-strains which provide the bases for the differential diagnosis.

### The predicted epitopes induce neutralization antibody production

SARS-CoV-2 pseudo-virus neutralization assay is a well-accepted method to detect the ability of vaccine to inhibit SARS-CoV-2 infection (Ni et al., 2020; Wang et al., 2020). To assess neutralization antibodies induced by S protein epitopes, we incubated the immunization sera with D614 or G614 SARS-CoV-2 pseudo-viruses and then the mixture was added to ACE2-293FT cells which stably expressed ACE2. The results showed that immunized sera of S92-106, S139-153, S439-454 and S455-469 epitopes inhibited SARS-CoV-2 pseudo-virus infection compared to HBc-S control (p < 0.0001), with inhibition rates around 40%-50% (Fig. 3A). Also, the sera of S16-30, S243-257, S406-420, S475-499, S556-570, S793-812(N) and S909-923 inhibited SARS-CoV-2 infection with the inhibition rate from 20% to 40% (Fig. 3A), indicating that these 11 epitopes induced neutralization antibody production. To detect the effect of epitope immunization on the neutralizing responses of G614 SARS-CoV-2, we incubated the epitope-immunized sera with the G614 SARS-CoV-2 pseudo-viruses. The results showed that sera of epitopes inhibiting D614 SARS-CoV-2 also inhibited G614 SARS-CoV-2 infection, except of S16-30, S243-257 and S556-570 (Fig. 3B). However, the immunized sera of epitopes S63-85, S495-509, S675-689, S703-719, S793-812, S1065-1079, S1065-1079(N), and S1106-1120 only inhibited G614 SARS-CoV-2 pseudo-virus infection. Interestingly, compared with its non-glycosylation epitope, S63-85(N), S703-719(N) and S1065-1079(N) induced less neutralizing antibodies to G614 pseudovirus while that of S793-812(N) increased (Fig. 3B). We then 2-fold serial diluted the sera with inhibition rate >50%, and determined the neutralizing antibody titers induced by these epitopes. S63-85 induced the highest neutralizing effect with antibody titer at 1:80 (Fig. 3C). The structural analysis showed that most of these neutralizing epitopes to D614 and G614 SARS-CoV-2 were in or near N-terminal domain (NTD), receptor-binding domain (RBD) or S2’ cleavage site of S protein and were spatially clustered (Fig. 3D-I), except S1106-1120 and S675-689 which are in or near transmembrane domain and S1/S2 cleavage site at interface of S1 and S2 subunits of S protein, respectively.

**Figure 3.**
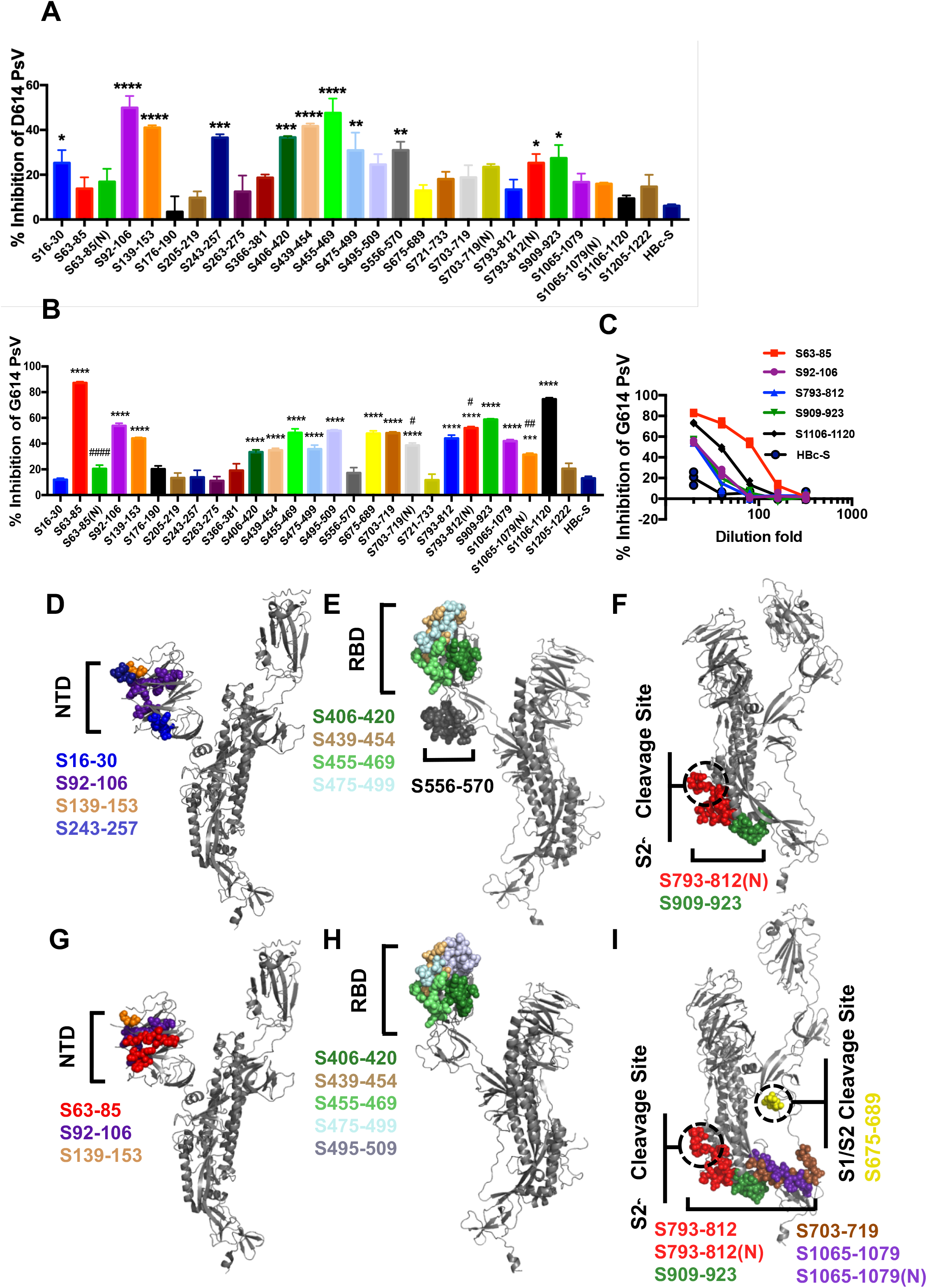
Antibodies induced by epitopes of protein S inhibit SARS-CoV-2 pseudo-virus infection. Pooled mice sera collected at day 10 after the third immunization were diluted (1:20) in DMEM, mixed with D614 (A) or G614 (B) SARS-CoV-2 pseudo-viruses (PsV) and incubated at 37 °C for 1 h. The mixture was then added to ACE2-293T. Firefly luciferase activity was measured 72 h post-infection (Compared with HBc-S control; *p < 0.05; **p < 0.01; ***p < 0.001; ****p < 0.0001; Compared with non-glycosylated epitope; ^#^p < 0.05; ^##^p < 0.01; ^####^p < 0.0001). (C) We 2-fold serial diluted the sera with inhibition rate >50% in DMEM, and mixed with SARS-CoV-2 pseudo-viruses. (D-I) Spatial positions of D614 (D-F) and G614 pseudovirus (G-I) neutralization epitopes (colored), respectively, in or near N-terminal domain (NTD) (D and G), receptor-binding domain (RBD) (E and H) and S2’ cleavage site (F and I) of S protein (grey).

## DISCUSSION

Vaccines are potent means to control the current pandemic of COVID-19 and to prevent future outbreak, thus fully understanding the immune responses elicited by the virus epitopes is urgent. As antigenic determinants, identifying and understanding epitopes would facilitate vaccine design and development. Since neutralization antibodies usually recognize the surface area of the virus proteins, identification of epitopes in surface area based on 3D structure of proteins may increase the efficiency to find the epitopes that elicit neutralization antibodies. In this study, we in first time used high-performance computer to simulate the three-dimensional structures of major proteins on SARS-CoV-2 and predicted 33 surface area epitopes using the modeled protein structures, which was proved to be efficient and accurate by the further mouse immunization and pseudo-virus neutralization assay. Within the 33 identified epitopes, 24 were conserved with >80% homology and 18 shared >90% homology among SARS-CoV-2, SARS-CoV and bat coronavirus RaTG13 (Table S1), implicating that these epitopes could be used as for designing broad-spectrum betacoronavirus vaccines.

Some surface area epitopes of SARS-CoV-2 were determined to be immunodominant in present study by detecting the binding of the antibodies in early convalescent sera of COVID-19 patient to various predicted epitopes. Consistent with previous report, S556-570 was an immunodominant epitope and this epitope was able to elicit neutralization antibodies (Fig. 3A) (Poh et al., 2020). S675-689 and S721-733 were immunodominant epitopes in imported strains but not in domestic strains, which may result from antigenic drift by the 614th amino acid variance on S protein of SARS-CoV-2 between imported and domestic cases. In most imported cases, glycine was in the 614th position of S protein, which possibly made the S675-689 and S721-733 regions more accessible by specifically destabilized the “up” state of the viral spike via unpacking with T859 in adjacent helical stalk (Becerra-Flores and Cardozo, 2020). Moreover, S556-570 was no longer an immunodominant and neutralizing epitope in the G614 strain and the antibodies induced by S675-689 inhibited G614 but not D614 pseudo-virus entry into ACE2 expressing 293FT cells, implicating an antigenic drift was caused by the D614G mutation. Two epitopes, N152-170 and N357-373, are highly conserved among the SARS-CoV-2, SARS-CoV and bat coronavirus RaTG13. Consistent with previous reports, these two epitopes were immunodominant sites (Guo et al., 2004). Importantly, they bound to neutralization antibodies in recovered SARS-CoV patients (Guo et al., 2004; Shichijo et al., 2004). M183-197 is another immunodominant epitope in domestic cases. Since this epitope is located in the S4 subsite of the active center of M protein, it is possible for it to elicit neutralization antibodies that inhibit the protease function of M protein (Dai et al., 2020; Yang et al., 2005). Interestingly, although no sequence variance was observed on M protein from all sequenced COVID-19 cases, two consecutive mutations (K203RR204G) were found in the highly conserved serine-rich linker region (LKR) of N protein. Since the LKR region is essential for flexibility of N protein and binds to M protein (Yang et al., 2005), it is possible that the R203G204 on N protein is relevant with the epitope immunodominance of M183-197. The difference of immunodominant epitopes from domestic and imported strains may have implications in designing assays for rapid classification and verification of virus sub-strains.

Eleven and seventeen epitope regions epitopes were found to elicit neutralization antibodies that inhibit the cell entry of D614 and G614 pseudo-virus, respectively. The antibodies induced by four epitopes (S406-420, S439-454, S455-469, and S475-499) but not S366-381 or S495-509 in RBD region exhibited neutralization effect on both D614 and G614 pseudo-virus, which is consistent with the interaction interface between SARS-CoV-2 receptor-binding motif (RBM) and ACE2 (Seydoux et al., 2020; Shang et al., 2020), indicating that these epitopes are suitable for designing universal vaccines. Previous report showed that a cryptic epitope in the trimeric interface of S protein induced neutralization antibodies for SARS-CoV but not SARS-CoV2. Consistently, the antibodies induced by epitope S366-381 did not show neutralization effect on the entry of the pseudo-virus (Wrapp et al., 2020; Yuan et al., 2020b). Interestingly, not only the antibodies targeting the interaction interface between RBD and ACE2, but also the antibodies binding with N-terminal domain (NTD) of S protein, such as S16-30, S92-106, S139-153 and S243-257 showed neutralization effect on D614 strain. Within the neutralizing NTD epitopes, S92-106 and S139-153 also showed neutralization effect on G614 strain. Antibodies induced by S63-85 but not its glycosylated form inhibited the cell entry of G614 pseudovirus rather than D614 pseudovirus, and the epitopes S703-719(N) and S1064-1079(N) induced less neutralizing antibodies compared to the unglycosylated ones, indicating native oligomannose and complex-type N-glycan might pose a shield effect on the epitope and mutation at the 614^th^ position possibly affected the exposure of the epitope by altering the pose of the glycan shield (Barnes et al., 2018). In contrast, the glycosylated epitope S793-812(N) showed more inhibitory effect than that of S793-812 on both G614 and D614 pseudoviruses, suggesting that glycosylation on the epitope might affect viral infectivity. Two conserved epitopes (S793-812(N) and S909-923) near the highly-conserved S2’ protease cleavage site of S protein also induced neutralization antibodies on both D614 and G614 pseudo-virus, indicating that there may be mechanism by which blocks cell entry of SARS-CoV-2. Notably, several identified neutralizing epitopes are consistent with the epitopes of some important neutralizing antibodies, such as S139-153 to antibody 4A8 (PDB 7C2L), S406-420 to antibody C105 (PDB 6XCN) and S16-30 to antibody P2B-2F6 (PDB 7BWJ) (Barnes et al., 2020; Chi et al., 2020; Ju et al., 2020), suggesting these epitopes might be the antibody-targeting sites. Importantly, we first found a shift of immunodominant and neutralizing epitope region from S556-S570 to S675-689 upon the D614G mutation. S675-689 is at the S1/S2 cleavage site located at interface of S1 and S2 subunits of S protein which is important for spike protein mediated virus-cell membrane fusion. Our results suggested that the S675-689 epitope was at the vulnerability site of SARS-CoV-2 and might be an ideal candidate and targeting site for vaccine development. Moreover, our results showed that the neutralizing epitopes are highly spatial clustered, indicating that conformational epitopes in the above regions may be used for designing an effective vaccine.

In conclusion, we have successfully predicted SARS-CoV-2 epitopes based on of the 3D structures of S, M, N, E proteins, validated their immunogenicity, characterized the homology of the epitopes among betacoronavirus, and identified the neutralization and immunodominant epitopes (Table S3). Our findings provide a wide neutralization and immunodominant epitope spectrum for the design of an effective, safe vaccine, differential diagnosis and virus classification.

## MATERIALS AND METHODS

### KEY RESOURCES TABLE

| REAGENT | SOURCE | IDENTIFIER |
| --- | --- | --- |
| Antibodies |  |  |
| HRP-conjugated goat anti-mouse IgG | Abcam | Cat# ab6789 |
| HRP-conjugated goat anti-human IgG | Abcam | Cat# ab6858 |
| Chemicals, Peptides, and Recombinant Proteins |  |  |
| BSA | Sigma-Aldrich Corporation | Cat# V900933 |
| Epitope peptides | This paper | N/A |
| HBc-S | This paper | N/A |
| Imject™ Alum Adjuvant | Thermo Fisher | Cat# 77161 |
| D614 SARS-CoV-2 pseudoviruses | PackGene Biotech | Cat# LV-nCov1 |
| G614 SARS-CoV-2 pseudoviruses | PackGene Biotech | Cat# LV-nCov1 |
| chromogenic substrate TMB | Thermo Fisher | Cat# 34028 |
| SARS-CoV-2 Spike S1+S2 | Sino Biological Inc. | Cat# 40589-V08B1 |
| SARS-CoV-2 nucleocapsid | Sino Biological Inc. | Cat#40588-V08B |
| Critical Commercial Assays |  |  |
| Bright-Glo™ Luciferase Assay System | Promega | Cat# E2620 |
| SARS-CoV-2 Nucleic Acid Extraction Kit | Daan Gene | Cat#DA0931 |
| Experimental Models: Cell Lines |  |  |
| ACE2-239T cells | PackGene<br>Biotech | Cat#nCov-3 |
| Software and Algorithms |  |  |
| Gromacs v5.1 | GROMACS | http://www.gromacs.org/ |
| Discotope 2.0 | Immune epitope<br>database and<br>analysis<br>resource (IEDB) | http://tools.iedb.org/discotope/ |
| ClustalW2 | EMBL's<br>European<br>Bioinformatics<br>Institute<br>(EMBL-EBI) | https://www.ebi.ac.uk/Tools/msa/clustalw2/ |

### EXPERIMENTAL MODEL AND SUBJECT DETAILS

#### Specimens from SARS-CoV-2 patients

Serum samples were collected from 20 early convalescent patients with COVID-19 which were confirmed by SARS-CoV-2 real-time reverse transcriptase–polymerase chain reaction (RT-PCR). 12 patients were infected in China and the other 8 were imported cases from Europe. The median age of imported and domestic patients was 50.8 years (range, 10-72 years) and 30.6 years (range, 17-50 years), respectively. This study was approved by the Institutional Review Board of the Fifth Hospital of Shijiazhuang. Waiver of informed consent for collection of samples from infected individuals was granted by the institutional ethics committee. Nucleic acids from throat swab samples were extracted using SARS-CoV-2 Nucleic Acids Extraction Kit (Daan Gene, Zhongshan, China) according to the manufacturer’s instructions. The genes of S, N, E and M were reverse transcripted, amplified and sequenced.

#### Mice

6-8 week-old BALB/c female mice were obtained from Beijing HFK Bioscience CO., LTD (Beijing, China) and maintained with access to food and water ad libitum in a colony room kept at 22 ± 2 °C and 50 ± 5% humidity, under a 12:12 light/dark cycle. All animal experiments were performed in accordance with the China Public Health Service Guide for the Care and Use of Laboratory Animals. Experiments involving mice and protocols were approved by the Institutional Animal Care and Use Committee of Tsinghua University (AP#15-LRT1).

### METHOD DETAILS

#### Epitope prediction

Homologous modeling and molecular dynamics simulation was used to predict the structure of S, M, N, E protein. The genome sequence of SARS-CoV-2 isolate Wuhan-Hu-1 was retrieved from the NCBI database under the accession number MN988669.1 and the protein sequences were acquired according to the annotation. The original pdb file of the S, M, N, E protein was established by homologous modeling using SWISS-MODEL (Waterhouse et al., 2018) according to the template structures of SARS-CoV spike glycoprotein (PDB: 6ACC), SARS-CoV main peptidase M(pro) (PDB: 2A5K), SARS-CoV envelope small membrane protein (PDB: 5X29) and SARS-CoV nucleocapsid protein (PDB: 1SSK), respectively. On the basis of the homologous modelled pdb file, added water, adjusted pH of chloride and sodium ions and ran molecular dynamics simulation program, obtaining the pdb file in the human body temperature (310K) state. We then calculated the full atomic structure of the protein for 1 μs using the molecular dynamics software GROMACS 5.1 on Sunway TaihuLight supercomputer and obtained the molecular orbital energy level and spatial structure information of the protein. In particular, the RBD region was referred to as the fragment from 347 to 520 amino acid of S protein. Structure-based conformational B cell epitope prediction was performed by using Discotope 2.0 (Kringelum et al., 2012) and −2.5 was used as a positivity cutoff. All the appropriate linear epitope windows were then selected by the following criteria: 1) solvate accessible regions with high surface probability; 2) regions with high antigenicity and flexibility; 3) the key functional residues of conformational B cell epitopes were centralized and with high degree of continuity in the window. The selected epitopes were then applied to homology analysis with the according sequences from SARS-CoV and RaTG13 via ClustalW (Thompson et al., 1994). N-glycosylated regions with homology > 50% were selected for synthesis of N-glycosylated epitopes. Epitope homology was calculated by the following formula:

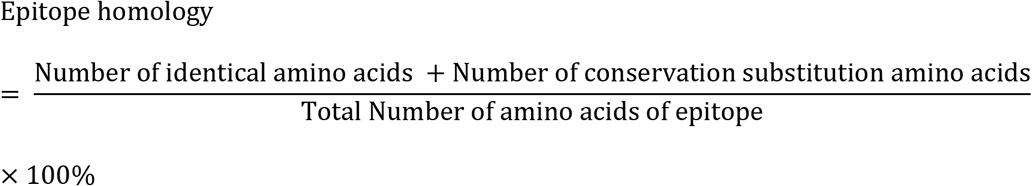

#### Preparation and characterization of HBc-S-peptide VLP vaccine

HBC-SpyCatcher (HBc-S) was purified as described previously (Ji et al., 2018). Purified protein was concentrated and determined by BCA protein assay kit (Pierce, Rockford, IL, USA). The purity of the recombinant protein was analyzed by SDS-PAGE. The peptides of SpyTag-epitope were chemically synthesized by GL Biochem (Shanghai, China). For the preparation of epitope conjugated HBc-S VLP vaccine, HBc-S VLPs were incubated with 3-fold molar excess of epitope peptide for 3 h at room temperature in citrate reaction buffer (40 mM Na_2_HPO_4_, 200 mM sodium citrate, pH 6.2), and was then dialyzed with 100 kDa cut-off membrane to remove the unreacted epitope peptide.

Transmission electron microscopy (TEM) was used for the morphological examination of HBc-S-peptide VLP vaccine. 20 μl VLPs (0.1-0.3 mg/ml) were applied to 200 mesh copper grids for 5 min and negatively stained with 2% uranyl acetate for 1 min, and then blotted with filter paper and air dried. VLPs were imaged in a Hitachi TEM system at 80 kV at 40,000 × magnification. To measure the hydrodynamic size of HBc-S-peptide VLP vaccine using dynamic light scattering (DLS), 9 μL of HBc-S or HBc-S-P VLPs at concentration of 0.1 mg/mL was loaded into a Uni tube and held at 2 min at room temperature. Each measurement was taken four times with 5 DLS acquisitions by an all-in-one stability platform Uncle (Unchained lab, USA).

#### Mice immunization

To evaluate the immunogenicity of the epitopes, female BALB/c mice (6-8 weeks) were subcutaneously vaccinated with HBc-S-P VLPs (containing 100 μg HBc-S) in citrate buffer (200 mM citrate acid, 40 mM NaH_2_PO_4_, pH 6.2, 100 μl) mixed with Alum (1:3 v/v) (ThermoFisher, Waltham, MA, USA). HBc-S was used as a control. Each group of mice (n=5) received their first injection at day 0 and boosters at day 14 and 28. Serum samples were taken 10 days after each immunization.

#### Enzyme-linked immunosorbent assay (ELISA)

Serum antibodies specific for epitope peptides and SARS-CoV-2 proteins were detected by ELISA. 96-well plates (Dynex Technologies, Chantilly, VA) were coated with 0.5 μg peptides, 100 ng S or N protein per well at 4°C overnight, respectively, and then washed 3 times with PBS and blocked with 3% BSA (in 0.1% PBST) for 2 h at 37 °C. After blocking, the plates were incubated with serial dilutions of the sera (100 μl/well, in two-fold dilution) for 2 h at 37 °C. The bound serum antibodies were detected with HRP-conjugated goat anti-mouse IgG (Zhongshan Golden Bridge Biotechnology Co., Beijing, China) and chromogenic substrate TMB (ThermoFisher, Waltham, MA, USA). The cut-off for seropositivity was set as the mean value plus three standard deviations (3SD) in HBc-S control sera. The binding of the epitopes to the sera of COVID-19 infected patients were detected by ELISA using the same procedure as described above, 96-well plates were coated with 0.5 μg peptides and sera were diluted at 1:50. The cut-off lines were based on the mean value + 3SD in 4-5 healthy persons. All ELISA studies were performed at least twice.

#### SARS-CoV-2 pseudovirus neutralization assay

Pooled mice sera collected at day 10 after the third immunization were diluted in DMEM supplemented with 10% fetal bovine serum, mixed with 1.6×10^6^ SARS-CoV-2 pseudoviruses and incubated at 37 °C for 1 h. The mixture was then added to 1.5×10^4^ ACE2-293T cells and the medium was replaced after 6 h. Firefly luciferase activity was measured 72 h post-infection using Bright-Glo™ Luciferase Assay System (Promega). All neutralization studies were performed at least twice. Three independently mixed replicates were measured for each experiment.

#### Statistical analysis

The data presented in this study were expressed as mean ± SEM. Data were analyzed by one-way (ANOVA), followed by multiple comparisons using Dunnett’s test within GraphPad Prism 7.0 software. Student t-test was used to analyze the data of non-glycosylated and glycosylated epitopes. p < 0.05 was considered to be significant.

## ACKNOWLEDGEMENTS

This work was supported by grants from the National Natural Science Foundation of China (81971610, 81971073 and 81903531), the National Science and Technology Major Projects of New Drugs (2018ZX09733001-001-008), Innovation Academy for Green Manufacture, Chinese Academy of Sciences (IAGM2020C29) and Zhejiang University special scientific research fund for COVID-19 prevention and control (2020XGZX075).

## AUTHOR CONTRIBUTIONS

R.-T.L. designed the experiment and wrote the manuscript; S.L. designed the experiment, obtained the 3D structures, predicted epitopes and wrote the manuscript; X.-X.X. designed the experiment, performed the ELISA, statistical analysis and wrote the manuscript; L.Z. collected the patient’s blood samples and carried out the ELISA experiment; B.W., J.Z. carried out the experimental works involving protein purification, DLS, TEM, mice immunization, ELISA and the neutralization assay. T.-R.Y. carried out protein purification, ELISA; M.J, C.-P. L carried out the neutralization assay. C.-G.J. helped to design the study. D.-Q.L., L.Z., S.-J.H. and X.-L.Y. participated in the mice blood collection and mice immunization. G.-W.Y, X.-M.F. helped to obtain the 3D protein structures and high performance computation. H.-X.G. and Y.-L.W. collected the patient’s throat swab samples and extracted nucleic acids. J.X. and X.-H.S. collected the patient’s blood samples. C.-W.K. and B.-X.K. helped to perform the neutralization assay.

