## supplement for "The immunodominant and neutralization linear epitopes for SARS-CoV-2"

**Supplemental information**

**Figure S1.** Homology of the predicted epitopes of S, M, E, N proteins among SARS-CoV-2, SARS-CoV and bat coronavirus RaTG13. Related to Figure 1.

### S Protein

```
SARS-CoV        MFIFLLFLTLTSGSDLDRCTTFDDVQAPNYTQHTSSMRGVYYPDEIFRSDTLYLTQDLFL
SARS-CoV-2      MFVFLVLLPLVSSQCVNLTT--RTQLPPAY--TNSFTRGVYYPDKVFRSSVLHSTQDLFL
RaTG13          MFVFLVLLPLVSSQCVNLTT--RTQLPPAY--TNSSTRGVYYPDKVFRSSVLHLTQDLFL
                **:**::* *.*.. ::  *       * *   .*  *******::***..*: ******

SARS-CoV        PFYSNVTGFHTIN-------HTFDNPVIPFKDGIYFAATEKSNVVRGWVFGSTMNNKSQS
SARS-CoV-2      PFFSNVTWFHAIHVSGTNGTKRFDNPVLPFNDGVYFASTEKSNIIRGWIFGTTLDSKTQS
RaTG13          PFFSNVTWFHAIHVSGTNGIKRFDNPVLPFNDGVYFASTEKSNIIRGWIFGTTLDSKTQS
                **:**** **:*:       : *****:**:**:***:*****::***:**:*::.*:**

SARS-CoV        VIIINNSTNVVIRACNFELCDNPFFAVSKPMGT----QTHTMIFDNAFNCTFEYISDAFS
SARS-CoV-2      LLIVNNATNVVIKVCEFQFCNDPFLGVYYHKNNKSWMESEFRVYSSANNCTFEYVSQPFL
RaTG13          LLIVNNATNVVIKVCEFQFCNDPFLGVYYHKNNKSWMESEFRVYSSANNCTFEYVSQPFL
                ::*:**:*****:.*:*::*::**:.*    ..    ::.  ::..* ******:*: *

SARS-CoV        LDVSEKSGNFKHLREFVFKNKDGFLYVYKGYQPIDVVRDLPSGFNTLKPIFKLPLGINIT
SARS-CoV-2      MDLEGKQGNFKNLREFVFKNIDGYFKIYSKHTPINLVRDLPQGFSALEPLVDLPIGINIT
RaTG13          MDLEGKQGNFKNLREFVFKNIDGYFKIYSKHTPINLVRDLPPGFSALEPLVDLPIGINIT
                :*:. *.****:******** **:: :*. : **::***** **.:*:*:..**:*****

SARS-CoV        NFRAILTAFL------PAQDTWGTSAAAYFVGYLKPTTFMLKYDENGTITDAVDCSQNPL
SARS-CoV-2      RFQTLLALHRSYLTPGDSSSGWTAGAAAYYVGYLQPRTFLLKYNENGTITDAVDCALDPL
RaTG13          RFQTLLALHRSYLTPGDSSSGWTAGAAAYYVGYLQPRTFLLKYNENGTITDAVDCALDPL
                .*:::*: .        :.. * :.****:****:* **:***:***********: :**

SARS-CoV        AELKCSVKSFEIDKGIYQTSNFRVVPSRDVVRFPNITNLCPFGEVFNATKFPSVYAWERK
SARS-CoV-2      SETKCTLKSFTVEKGIYQTSNFRVQPTESIVRFPNITNLCPFGEVFNATRFASVYAWNRK
RaTG13          SETKCTLKSFTVEKGIYQTSNFRVQPTDSIVRFPNITNLCPFGEVFNATTFASVYAWNRK
                :* **::*** ::*********** *: .:******************* * *****:**

SARS-CoV        RISNCVADYSVLYNSTFFSTFKCYGVSATKLNDLCFSNVYADSFVVKGDDVRQIAPGQTG
SARS-CoV-2      RISNCVADYSVLYNSASFSTFKCYGVSPTKLNDLCFTNVYADSFVIRGDEVRQIAPGQTG
RaTG13          RISNCVADYSVLYNSTSFSTFKCYGVSPTKLNDLCFTNVYADSFVITGDEVRQIAPGQTG
                ***************: ********** ********:********: **:**********

SARS-CoV        VIADYNYKLPDDFMGCVLAWNTRNIDATSTGNYNYKYRYLRHGKLRPFERDISNVPFSPD
SARS-CoV-2      KIADYNYKLPDDFTGCVIAWNSNNLDSKVGGNYNYLYRLFRKSNLKPFERDISTEIYQAG
RaTG13          KIADYNYKLPDDFTGCVIAWNSKHIDAKEGGNFNYLYRLFRKANLKPFERDISTEIYQAG
                 ************ ***:***:.::*:.  **:** ** :*:.:*:*******.  :. .

SARS-CoV        GKPCTP-PALNCYWPLNDYGFYTTTGIGYQPYRVVVLSYELLNAPATVCGPKLSTDLIKN
SARS-CoV-2      STPCNGVEGFNCYFPLQSYGFQPTNGVGYQPYRVVVLSFELLHAPATVCGPKKSTNLVKN
RaTG13          SKPCNGQTGLNCYYPLYRYGFYPTDGVGHQPYRVVVLSFELLNAPATVCGPKKSTNLVKN
                ..**.   .:***:**  ***  * *:*:*********:***:********* **:*:**

SARS-CoV        QCVNFNFNGLTGTGVLTPSSKRFQPFQQFGRDVSDFTDSVRDPKTSEILDISPCSFGGVS
SARS-CoV-2      KCVNFNFNGLTGTGVLTESNKKFLPFQQFGRDIADTTDAVRDPQTLEILDITPCSFGGVS
RaTG13          KCVNFNFNGLTGTGVLTESNKKFLPFQQFGRDIADTTDAVRDPQTLEILDITPCSFGGVS
                :**************** *.*:* ********::* **:****:* *****:********

SARS-CoV        VITPGTNASSEVAVLYQDVNCTDVSTAIHADQLTPAWRIYSTGNNVFQTQAGCLIGAEHV
SARS-CoV-2      VITPGTNTSNQVAVLYQDVNCTEVPVAIHADQLTPTWRVYSTGSNVFQTRAGCLIGAEHV
RaTG13          VITPGTNASNQVAVLYQDVNCTEVPVAIHADQLTPTWRVYSTGSNVFQTRAGCLIGAEHV
                *******:*.:***********:* .*********:**:****.*****:**********
```

```
SARS-CoV        DTSYECDIPIGAGICASYHTVSL----LRSTSQKSIVAYTMSLGADSSIAYSNNTIAIPT
SARS-CoV-2      NNSYECDIPIGAGICASYQTQTNSPRRARSVASQSIIAYTMSLGAENSVAYSNNSIAIPT
RaTG13          NNSYECDIPIGAGICASYQTQTNS----RSVASQSIIAYTMSLGAENSVAYSNNSIAIPT
                :.****************:* :      **.:.:**:********:.*:*****:*****

SARS-CoV        NFSISITTEVMPVSMAKTSVDCNMYICGDSTECANLLLQYGSFCTQLNRALSGIAAEQDR
SARS-CoV-2      NFTISVTTEILPVSMTKTSVDCTMYICGDSTECSNLLLQYGSFCTQLNRALTGIAVEQDK
RaTG13          NFTISVTTEILPVSMTKTSVDCTMYICGDSTECSNLLLQYGSFCTQLNRALTGIAVEQDK
                **:**:***::****:******.**********:*****************:***.***:

SARS-CoV        NTREVFAQVKQMYKTPTLKDFGGFNFSQILPDPLKSTKRSFIEDLLFNKVTLADAGFMKQ
SARS-CoV-2      NTQEVFAQVKQIYKTPPIKDFGGFNFSQILPDPSKPSKRSFIEDLLFNKVTLADAGFIKQ
RaTG13          NTQEVFAQVKQIYKTPPIKDFGGFNFSQILPDPSKPSKRSFIEDLLFNKVTLADAGFIKQ
                **:********:**** :*************** * :********************:**

SARS-CoV        YGECLGDINARDLICAQKFNGLTVLPPLLTDDMIAAYTAALVSGTATAGWTFGAGAALQI
SARS-CoV-2      YGDCLGDIAARDLICAQKFNGLTVLPPLLTDEMIAQYTSALLAGTITSGWTFGAGAALQI
RaTG13          YGDCLGDIAARDLICAQKFNGLTVLPPLLTDEMIAQYTSALLAGTITSGWTFGAGAALQI
                **:***** **********************:*** **:**::** *:************

SARS-CoV        PFAMQMAYRFNGIGVTQNVLYENQKQIANQFNKAISQIQESLTTTSTALGKLQDVVNQNA
SARS-CoV-2      PFAMQMAYRFNGIGVTQNVLYENQKLIANQFNSAIGKIQDSLSSTASALGKLQDVVNQNA
RaTG13          PFAMQMAYRFNGIGVTQNVLYENQKLIANQFNSAIGKIQDSLSSTASALGKLQDVVNQNA
                ************************* ******.**.:**:**::*::*************

SARS-CoV        QALNTLVKQLSSNFGAISSVLNDILSRLDKVEAEVQIDRLITGRLQSLQTYVTQQLIRAA
SARS-CoV-2      QALNTLVKQLSSNFGAISSVLNDILSRLDKVEAEVQIDRLITGRLQSLQTYVTQQLIRAA
RaTG13          QALNTLVKQLSSNFGAISSVLNDILSRLDKVEAEVQIDRLITGRLQSLQTYVTQQLIRAA
                ************************************************************

SARS-CoV        EIRASANLAATKMSECVLGQSKRVDFCGKGYHLMSFPQAAPHGVVFLHVTYVPSQERNFT
SARS-CoV-2      EIRASANLAATKMSECVLGQSKRVDFCGKGYHLMSFPQSAPHGVVFLHVTYVPAQEKNFT
RaTG13          EIRASANLAATKMSECVLGQSKRVDFCGKGYHLMSFPQSAPHGVVFLHVTYVPAQEKNFT
                **************************************:**************:**:***

SARS-CoV        TAPAICHEGKAYFPREGVFVFNGTSWFITQRNFFSPQIITTDNTFVSGNCDVVIGIINNT
SARS-CoV-2      TAPAICHDGKAHFPREGVFVSNGTHWFVTQRNFYEPQIITTDNTFVSGNCDVVIGIVNNT
RaTG13          TAPAICHDGKAHFPREGVFVSNGTHWFVTQRNFYEPQIITTDNTFVSGSCDVVIGIVNNT
                *******:***:******** *** **:*****:.*************.*******:***

SARS-CoV        VYDPLQPELDSFKEELDKYFKNHTSPDVDLGDISGINASVVNIQKEIDRLNEVAKNLNES
SARS-CoV-2      VYDPLQPELDSFKEELDKYFKNHTSPDVDLGDISGINASVVNIQKEIDRLNEVAKNLNES
RaTG13          VYDPLQPELDSFKEELDKYFKNHTSPDVDLGDISGINASVVNIQKEIDRLNEVAKNLNES
                ************************************************************

SARS-CoV        LIDLQELGKYEQYIKWPWYVWLGFIAGLIAIVMVTILLCCMTSCCSCLKGACSCGSCCKF
SARS-CoV-2      LIDLQELGKYEQYIKWPWYIWLGFIAGLIAIVMVTIMLCCMTSCCSCLKGCCSCGSCCKF
RaTG13          LIDLQELGKYEQYIKWPWYIWLGFIAGLIAIIMVTIMLCCMTSCCSCLKGCCSCGSCCKF
                *******************:***********:****:*************.*********

SARS-CoV        DEDDSEPVLKGVKLHYT     1255
SARS-CoV-2      DEDDSEPVLKGVKLHYT     1273
RaTG13          DEDDSEPVLKGVKLHYT     1269
                *****************
```

### M Protein

```
SARS-CoV        -MADNGTITVEELKQLLEQWNLVIGFLFLAWIMLLQFAYSNRNRFLYIIKLVFLWLLWPV
SARS-CoV-2      MADSNGTITVEELKKLLEQWNLVIGFLFLTWICLLQFAYANRNRFLYIIKLIFLWLLWPV
RaTG13          -MADNGTITVEELKKLLEQWNLVIGFLFLTWICLLQFAYANRNRFLYIIKLIFLWLLWPV
                   .**********:**************:** ******:***********:********

SARS-CoV        TLACFVLAAVYRINWVTGGIAIAMACIVGLMWLSYFVASFRLFARTRSMWSFNPETNILL
SARS-CoV-2      TLACFVLAAVYRINWITGGIAIAMACLVGLMWLSYFIASFRLFARTRSMWSFNPETNILL
RaTG13          TLACFVLAAVYRINWITGGIAIAMACLVGLMWLSYFIASFRLFARTRSMWSFNPETNILL
                ***************:**********:*********:***********************
```

```
SARS-CoV        NVPLRGTIVTRPLMESELVIGAVIIRGHLRMAGHSLGRCDIKDLPKEITVATSRTLSYYK
SARS-CoV-2      NVPLHGTILTRPLLESELVIGAVILRGHLRIAGHHLGRCDIKDLPKEITVATSRTLSYYK
RaTG13          NVPLHGTILTRPLLESELVIGAVILRGHLRIAGHHLGRCDIKDLPKEITVATSRTLSYYK
                ****:***:****:**********:*****:*** *************************

SARS-CoV        LGASQRVGTDSGFAAYNRYRIGNYKLNTDHAGSNDNIALLVQ
SARS-CoV-2      LGASQRVAGDSGFAAYSRYRIGNYKLNTDHSSSSDNIALLVQ
RaTG13          LGASQRVAGDSGFAAYSRYRIGNYKLNTDHSSSSDNIALLVQ
                *******. *******.*************:.*.********
```

### E protein

```
SARS-CoV-2      MYSFVSEETGTLIVNSVLLFLAFVVFLLVTLAILTALRLCAYCCNIVNVSLVKPSFYVYS
RaTG13          MYSFVSEETGTLIVNSVLLFLAFVVFLLVTLAILTALRLCAYCCNIVNVSLVKPSFYVYS
SARS-CoV        MYSFVSEETGTLIVNSVLLFLAFVVFLLVTLAILTALRLCAYCCNIVNVSLVKPTVYVYS
                ******************************************************:.****

SARS-CoV-2      RVKNLNSSR-VPDLLV
RaTG13          RVKNLNSSR-VPDLLV
SARS-CoV        RVKNLNSSEGVPDLLV
                ********. ******
```

### N Protein

```
SARS-CoV-2      MSDNGPQ-NQRNAPRITFGGPSDSTGSNQNGERSGARSKQRRPQGLPNNTASWFTALTQH
RaTG13          MSDNGPQ-NQRNAPRITFGGPSDSTGSNQNGERSGARPKQRRPQGLPNNTASWFTALTQH
SARS-CoV        MSDNGPQSNQRSAPRITFGGPTDSTDNNQNGGRNGARPKQRRPQGLPNNTASWFTALTQH
                ******* ***.*********:***..**** *.*** **********************

SARS-CoV-2      GKEDLKFPRGQGVPINTNSSPDDQIGYYRRATRRIRGGDGKMKDLSPRWYFYYLGTGPEA
RaTG13          GKEDLKFPRGQGVPINTNSSPDDQIGYYRRATRRIRGGDGKMKDLSPRWYFYYLGTGPEA
SARS-CoV        GKEELRFPRGQGVPINTNSGPDDQIGYYRRATRRVRGGDGKMKELSPRWYFYYLGTGPEA
                ***:*:*************.**************:********:****************

SARS-CoV-2      GLPYGANKDGIIWVATEGALNTPKDHIGTRNPANNAAIVLQLPQGTTLPKGFYAEGSRGG
RaTG13          GLPYGANKDGIIWVATEGALNTPKDHIGTRNPANNAAIVLQLPQGTTLPKGFYAEGSRGG
SARS-CoV        SLPYGANKEGIVWVATEGALNTPKDHIGTRNPNNNAATVLQLPQGTTLPKGFYAEGSRGG
                .*******:**:******************** **** **********************

SARS-CoV-2      SQASSRSSSRSRNSSRNSTPGSSRGTSPARMAGNGGDAALALLLLDRLNQLESKMSGKGQ
RaTG13          SQASSRSSSRSRNSSRNSTPGSSRGTSPARMAGNGSDAALALLLLDRLNQLESKMSGKGQ
SARS-CoV        SQASSRSSSRSRGNSRNSTPGSSRGNSPARMASGGGETALALLLLDRLNQLESKVSGKGQ
                ************..***********.******..*.::****************:*****

SARS-CoV-2      QQQGQTVTKKSAAEASKKPRQKRTATKAYNVTQAFGRRGPEQTQGNFGDQELIRQGTDYK
RaTG13          QQQSQTVTKKSAAEASKKPRQKRTATKQYNVTQAFGRRGPEQTQGNFGDQELIRQGTDYK
SARS-CoV        QQQGQTVTKKSAAEASKKPRQKRTATKQYNVTQAFGRRGPEQTQGNFGDQDLIRQGTDYK
                ***.*********************** **********************:*********

SARS-CoV-2      HWPQIAQFAPSASAFFGMSRIGMEVTPSGTWLTYTGAIKLDDKDPNFKDQVILLNKHIDA
RaTG13          HWPQIAQFAPSASAFFGMSRIGMEVTPSGTWLTYTGAIKLDDKDPNFKDQVILLNKHIDA
SARS-CoV        HWPQIAQFAPSASAFFGMSRIGMEVTPSGTWLTYHGAIKLDDKDPQFKDNVILLNKHIDA
                ********************************** **********:***:**********

SARS-CoV-2      YKTFPPTEPKKDKKKKADETQALPQRQKKQQTVTLLPAADLDDFSKQLQQSMSSADSTQA
RaTG13          YKTFPPTEPKKDKKKKADETQALPQRQKKQQTVTLLPAADLDDFSKQLQQSMSSADSTQA
SARS-CoV        YKTFPPTEPKKDKKKKTDEAQPLPQRQKKQPTVTLLPAADMDDFSRQLQNSMSGASADST
                ****************:**:* ******** *********:****:***:***.*.: .:

SARS-CoV-2      --
RaTG13          --
SARS-CoV        QA
```

**Figure S2.** Preparation of HBc-S VLPs displayed with the epitopes. Related to Figure 1.

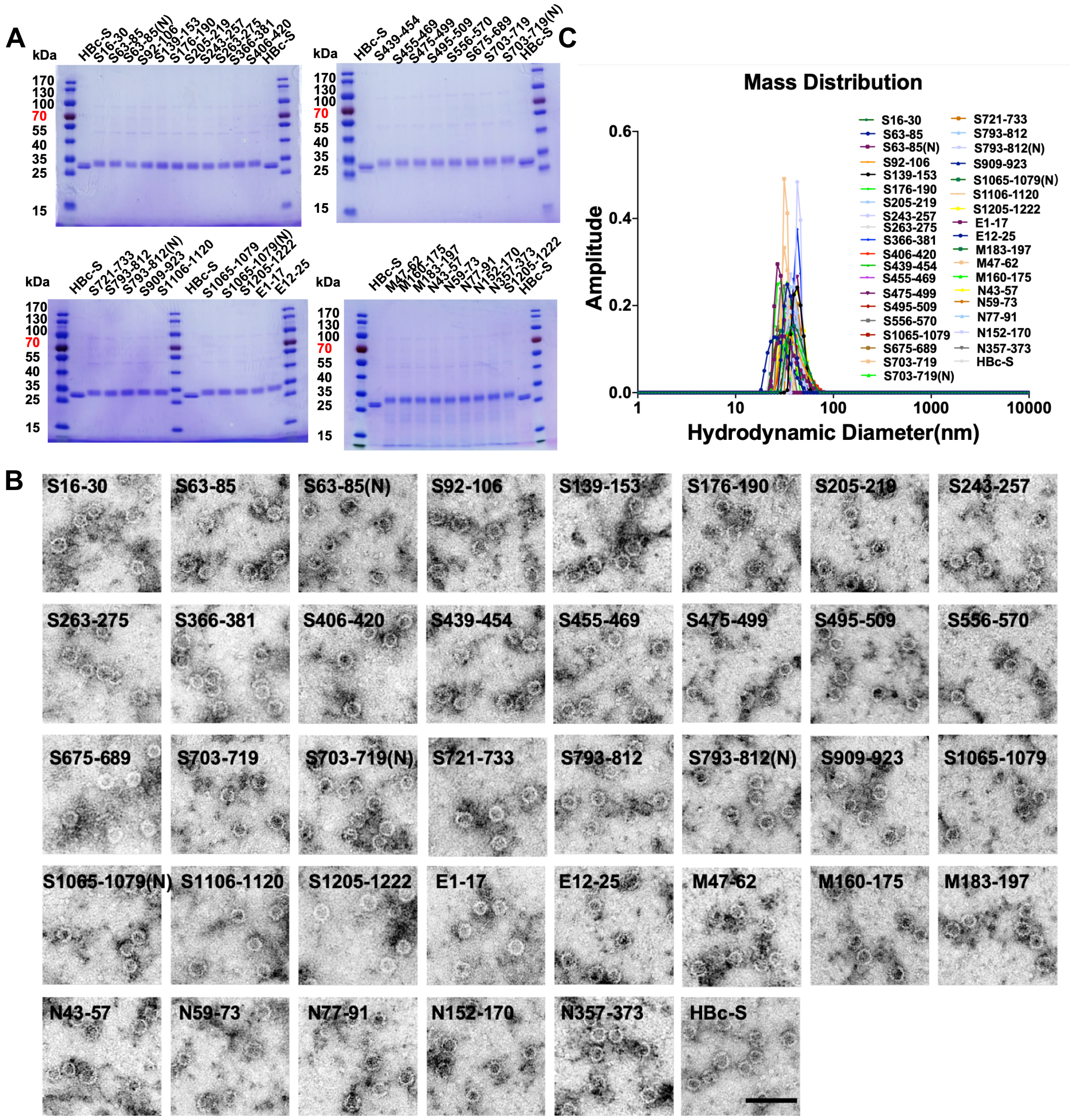


**Figure S2.** (A) SDS-PAGE analysis of the conjugation of HBc-S with epitope peptides. The lower band of HBc monomer is the cleavage product. (B) The morphology of HBc-S-P VLPs. HBc-S-P VLPs were imaged by Hitachi TEM at 80 KV at 40,000× magnification, the scale bar is 200 nm. (C) The representative hydrodynamic diameter of HBc-S-P VLPs.

**Figure S3.** Epitope conjugated on HBc-S VLPs induced high antibody titers against epitope peptides and S, N proteins at 10 days after the second immunization. Related to Figure 1.

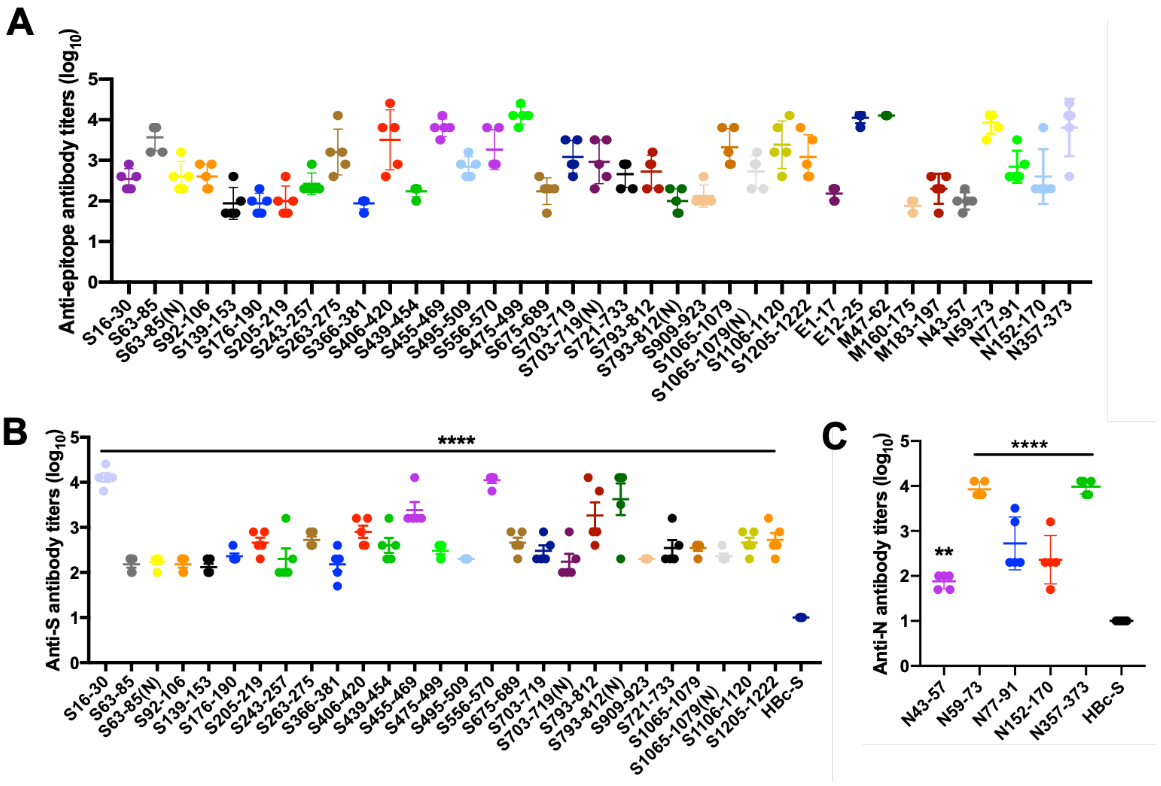

**Figure S4.** The non-immunodominant epitopes against S, E, M, N in early convalescent sera from COVID-19 patients. Related to Figure 2.

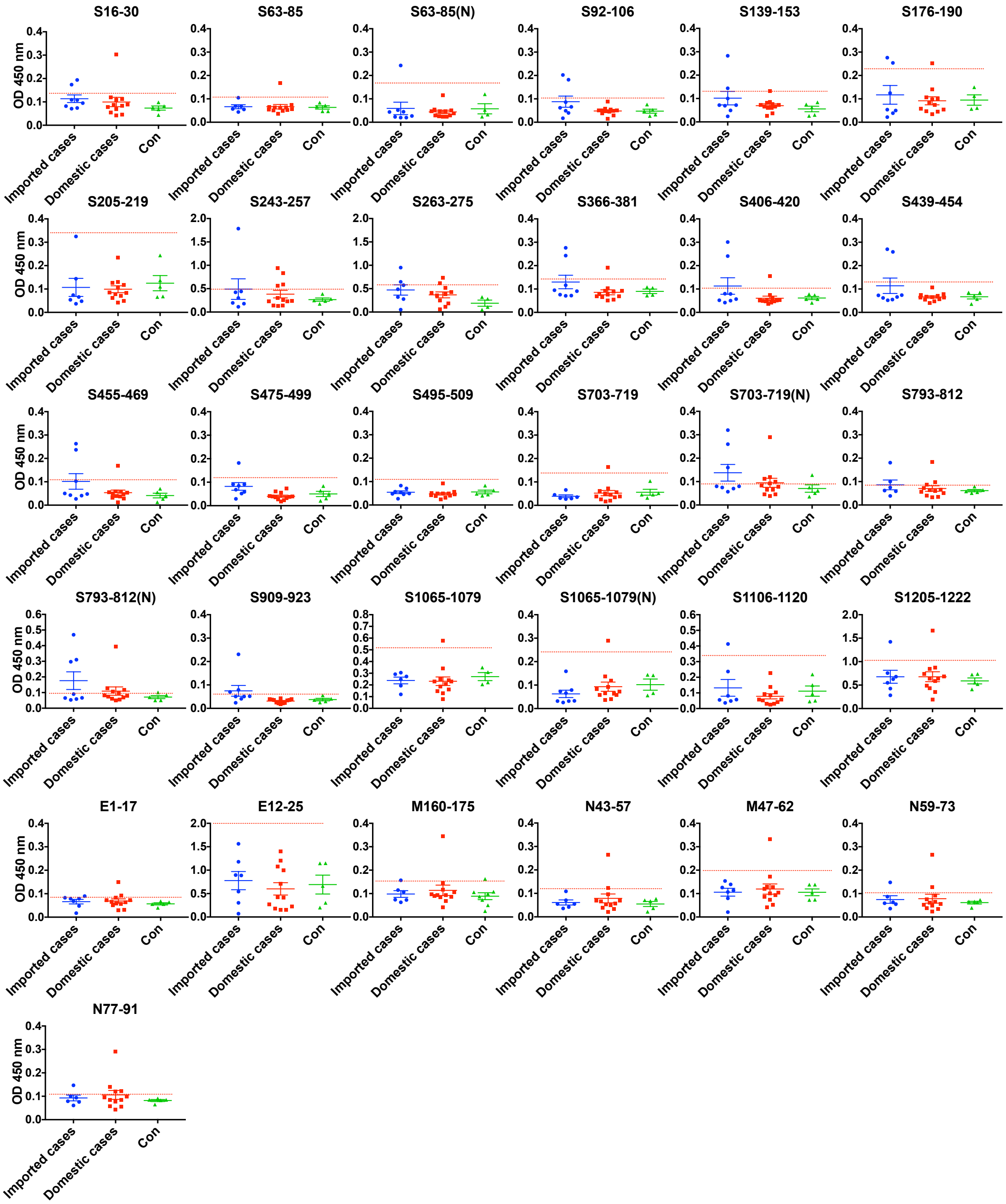

**Table S1.** Predicted epitope peptides. Related to Figure 1.

| Location | Sequence | Homology |
|---|---|---|
| S16-30 | VNLTTRTQLPPAYTN | 33.30% |
| S63-85 | TWFHAIHVSGTNGTKRFDNPVLP | 60.90% |
| S63-85(N) | TWFHAIHVSGTNGTKRFDN(GlcNAc)PVLP | 60.90% |
| S92-106 | FASTEKSNIIRGWIF | 100% |
| S139-153 | PFLGVYYHKNNKSWM | 26.70% |
| S176-190 | LMDLEGKQGNFKNLR | 73.30% |
| S205-219 | SKHTPINLVRDLPQG | 73.30% |
| S243-257 | ALHRSYLTPGDSSSG | 13.30% |
| S263-275 | AAYYVGYLQPRTF | 92.30% |
| S366-381 | SVLYNSASFSTFKCYG | 93.70% |
| S406-420 | EVRQIAPGQTGKIAD | 93.30% |
| S439-454 | NNLDSKVGGNYNYLYR | 62.50% |
| S455-469 | LFRKSNLKPFERDIS | 86.70% |
| S475-499 | AGSTPCNGVEGFNCYFPLQSYGFQP | 48% |
| S495-509 | YGFQPTNGVGYQPYR | 80% |
| S556-570 | NKKFLPFQQFGRDIA | 86.70% |
| S675-689 | QTQTNSPRRARSVAS | 40% |
| S703-719 | NSVAYSNNSIAIPTNFT | 94.10% |
| S703-719(N) | NSVAYSNNSIAIPTN(GlcNAc)FT | 94.10% |
| S721-733 | SVTTEILPVSMTK | 100% |
| S793-812 | PIKDFGGFNFSQILPDPSKP | 85% |
| S793-812(N) | PIKDFGGFN(GlcNAc)FSQILPDPSKP | 85% |
| S909-923 | IGVTQNVLYENQKLI | 93.30% |
| S1065-1079 | VTYVPAQEKNFTTAP | 100% |
| S1065-1079(N) | VTYVPAQEKN(GlcNAc)FTTAP | 100% |
| S1106-1120 | QRNFYEPQIITTDNT | 93.30% |
| S1205-1222 | KYEQYIKWPWYIWLGFIA | 100% |
| E1-17 | MYSFVSEETGTLIVNSV | 100% |
| E12-25 | LIVNSVLLFLAFVV | 100% |
| M47-62 | YIIKLIFLWLLWPVTL | 100% |
| M160-175 | DIKDLPKEITVATSRT | 100% |
| M183-197 | ASQRVAGDSGFAAYS | 86.70% |
| N43-57 | QGLPNNTASWFTALT | 100% |
| N59-73 | HGKEDLKFPRGQGVP | 100% |
| N77-91 | NSSPDDQIGYYRRAT | 93.30% |
| N152-170 | ANNAAIVLQLPQGTTLPKG | 89.40% |
| N357-373 | IDAYKTFPPTEPKKDKK | 100% |

**Table S2.** The Hydrodynamic Diameter and PDI of HBc-S-P VLPs. Related to Figure 1.

| Sample | S16-30 | S63-85 | S63-85(N) | S92-106 | S139-153 | S176-190 | S205-219 | S243-257 | S263-275 | S366-381 | S406-420 | S439-454 | S455-469 |
|---|---|---|---|---|---|---|---|---|---|---|---|---|---|
| PDI | 0.08 | 0.07 | 0.18 | 0.19 | 0.04 | 0.06 | 0.05 | 0.01 | 0.04 | 0.01 | 0.01 | 0.01 | 0.06 |
| Hydrodynamic Diameter(nm) | 48.0±13.0 | 37.3±10.0 | 39.8±13.0 | 45.5±14.5 | 45.0±8.5 | 45.1±10.7 | 44.7±10.2 | 46.7±4.7 | 44.2±8.7 | 45.9±12.2 | 45.7±3.4 | 44.8±10.8 | 45.4±3.8 |
| **Sample** | **S475-499** | **S495-509** | **S556-570** | **S675-689** | **S703-719** | **S703-719(N)** | **S721-733** | **S793-812** | **S793-812(N)** | **S909-923** | **S1065-1079** | **S1065-1079(N)** | **S1106-1120** |
| PDI | 0.02 | 0.21 | 0.04 | 0.03 | 0.10 | 0.09 | 0.03 | 0.06 | 0.14 | 0.04 | 0.06 | 0.04 | 0.06 |
| Hydrodynamic Diameter(nm) | 44.8±9.1 | 45.5±10.8 | 46.6±8.8 | 45.6±8.3 | 34.9±11.2 | 35.4±11.0 | 45.2±7.9 | 37.9±9.5 | 41.6±15.3 | 44.8±9.0 | 37.2±8.8 | 37.9±9.5 | 48.0±12.1 |
| **Sample** | **S1205-1222** | **E1-17** | **E12-25** | **M47-62** | **M160-175** | **M183-197** | **N43-57** | **N59-73** | **N77-91** | **N152-170** | **N357-373** | **HBc-S** | |
| PDI | 0.06 | 0.12 | 0.08 | 0.15 | 0.01 | 0.02 | 0.10 | 0.14 | 0.01 | 0.04 | 0.04 | 0.06 | |
| Hydrodynamic Diameter(nm) | 45.1±11.4 | 35.4±13.1 | 34.1±9.3 | 48.0±10.5 | 37.1±3.7 | 44.3±6.1 | 39.4±12.6 | 39.3±7.4 | 44.7±4.7 | 43.8±6.0 | 36.7±7.4 | 36.7±5.2 | |

**Table S3. Characteristics of epitopes. Related to Figure 1-3.**

| Epitopes | S protein | | N protein | M protein | E protein |
|---|---|---|---|---|---|
| | non-glycosylation | Glycosylation | | | |
| Number | 23 | 4 | 5 | 3 | 2 |
| Conserved (>80% homology) | S92-106, S263-275, S366-381, S406-420, S439-454, S455-469, S495-509, S556-570, S703-719, S721-733, S793-812, S909-923, S1065-1079, S1106-1120 | S703-719(N), S793-812(N), S1065-1079(N) | N43-57, N59-73, N77-91, N152-170, N357-373 | M47-62, M160-175, M183-197 | E12-25, E55-69 |
| High immunogenicity (antibody titer>$10^4$) | S16-30, S205-219, S455-469, S475-499, S556-570, S721-733, S793-812, S1106-1120, S1205-1222 | S793-812(N) | N59-73, N353-373 | M47-62 | E12-25 |
| Immunodominant | | | | | |
| Imported cases* | S675-689, S721-733 | None | N152-170 | None | None |
| Domestic cases* | S556-570 | None | N357-373, N152-170 | M183-197 | None |
| Neutralizing | | | | | |
| D614 SARS-CoV-2 | S16-30, S92-106, S139-153, S243-275, S406-420, S439-454, S455-469, S475-499, S556-570, S909-923 | S793-812(N) | NT | NT | NT |
| G614 SARS-CoV-2 | S63-85, S92-106, S139-153, S406-420, S439-454, S455-469, S475-499, S495-509 , S675-689,S703-719, S793-812, S909-923, S1065-1079, S1106-1120 | S703-719(N), S793-812(N), S1065-1079(N) | NT | NT | NT |

***imported (Europe) cases infected SARS-CoV-2 in early April, 2020 were and domestic (China) cases in early February, 2020**

**NT represents not tested**
